# Distinct genetic architecture for trait mean and plasticity in chickpea

**DOI:** 10.1101/2024.12.10.625655

**Authors:** Ganesh Alagarasan, Rajeev K. Varshney, Eswarayya Ramireddy

**Author notes:** First and Co-corresponding author.

## Abstract

Crop plants often face rapid environmental changes, and their ability to adapt to novel conditions depends on whether specific traits can adjust to match these changes. This capacity for adjustment, known as phenotypic plasticity, is frequently regarded as a mere “response” rather than a distinct trait with its own genetic basis, particularly in agricultural science. As a result, plasticity is often oversimplified and viewed as an undesirable phenomenon when the goal is to achieve trait stability. In this study, we present a quantitative metric to measure phenotypic plasticity and provide evidence, through genome-wide association studies (GWAS) in chickpea, that plasticity is indeed a genetically based trait, comparable to any other agronomic trait. This foundational study paves the way for future GWAS efforts to harness phenotypic plasticity, offering new insights for crop improvement strategies.

## Main

Given the rapid climate change, a broader view on the genetics of crop adaptation such as phenotypic plasticity warrants closer examination. However, progress in this area has been limited. This is partly due to conceptual differences regarding plasticity across scientific disciplines and a predominant focus on trait stability under agricultural field conditions.

Historically, early biologists such as Lamarck (1809), Darwin (1859), and Weismann (1893) focused on whether phenotypic changes due to environmental influences—manifestations of plasticity—could be inherited. While Darwin did consider the heritability of the capacity for variation, the explicit exploration of plasticity as an inherent trait was not fully developed. Waddington (1953) came closest to this by demonstrating that plastic responses could become fixed in the genome through genetic assimilation, showing a direct inheritance mechanism for plasticity.

Following the advent of Mendelian genetics, which emphasized fixed inheritance patterns, the complexity of plasticity was often underrepresented. The evolution of plasticity has since sparked significant debates among scientists about the genetic factors contributing to this trait (Schlichting and Pigliucci, 1993; Via, 1993; Nicoglou, 2015). Building on this discourse, studies have further explored the genetic intricacies underlying plasticity across various species. For instance, Scheiner and Lyman (1991) studied *Drosophila melanogaster*, providing insights into plasticity as both an inherent trait and a product of gene-environment interactions. They demonstrated significant heritability for thorax size plasticity, indicating a genetic component. However, genetic correlations between thorax size and plasticity complicated the conclusion that plasticity is an independent trait with distinct genetics from the trait mean. Similarly, in *Arabidopsis thaliana*, heritability for plasticity was observed, but significant non-additive genetic effects (dominance and overdominance) indicated complex genetic control (Andreou et al., 2023). This complexity challenges the consideration of plasticity as a simple, distinct trait separate from other genetically influenced traits. Additionally, a long-term study of great tits (*Parus major*) in a Dutch population revealed significant heritable variation in the plasticity of laying dates (Nussey et al., 2005). However, this plasticity arises from complex gene-environment interactions. The study found a significant correlation between average laying date and plastic response to temperature, suggesting shared genetic architecture.

Scheiner suggested three genetic models of plasticity: overdominance, epistasis, and pleiotropy (Scheiner, 1993). These models, while foundational, have evolved with further understanding. In molecular biology, we now know that changes at every level—ranging from epigenetic modifications, transcriptional and translational regulation, post-translational modifications (PTMs), to protein interactions—can all be influenced by the environment. These changes result in new phenotype (Stern et al., 2007; Qian et al., 2011; Zhang et al., 2012; Chen et al., 2017; Domnauer et al., 2021; Steward et al., 2022; Leung et al., 2023; Battle et al., 2024). It is now a matter of interest to document which mechanisms are changing. It is important to distinguish between genetic/non-genetic effects that merely alter the phenotype and those that specifically contribute to plasticity as a trait. The genetics of plasticity should focus on how genetic variation enables organisms to exhibit variable phenotypes under different environmental conditions, rather than just the phenotypic changes themselves. Therefore, while genetic interactions such as pleiotropic and epistatic effects can shape phenotypic outcomes, they should not be viewed as the genetics of plasticity unless they specifically affect the capacity for plasticity.

Resistance to accepting plasticity as a trait largely stems from the lack of studies explicitly identifying genes controlling the plasticity of specific traits. Recently, the exploration of plasticity as a trait in agricultural science has been emerging. For instance, research by Tetard-Jones et al. (2011) in barley inbred lines, Li et al. (2018) in sorghum inbred lines, Kusmec et al. (2017), and Jin et al. (2023) in maize inbred lines, as well as Chen et al. (2024) in rice hybrid lines, has identified separate genes for trait mean and trait plasticity, each with different measures of plasticity. However, proving that genes for plasticity and mean are distinct in artificial populations has several drawbacks. These populations often have limited genetic diversity, restricting the generalizability of findings to broader populations. Non-random mating patterns and specific crossing schemes can introduce biases, complicating genetic association interpretations. Intentional selection for specific traits can skew the identification of relevant genes, and simplified gene interactions in these populations may not capture the complexity of natural settings. Therefore, while artificial populations provide valuable insights, complementing these studies with investigations in more genetically diverse and environmentally variable natural populations is essential for a comprehensive understanding.

To address this gap, we sought to investigate whether plasticity can be considered a separate trait with its own genetic architecture in crop plants. To quantify plasticity in our study, we used the following formula:

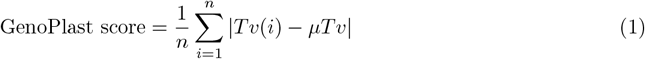

where:

- *Tv*(*i*): the trait value for a particular condition *i*.
- *µTv*: the median trait value across all conditions.
- |*Tv*(*i*) −*µTv* |: the absolute difference between the trait value for a specific condition and the median trait value. The absolute value ensures that the differences are considered without regard to their direction (positive or negative).
- *n*: the total number of conditions.

We then performed a genome-wide association study (GWAS) to identify the genetic loci associated with phenotypic plasticity in GAPIT 3.5 version (Wang and Zhang, 2021). The study employed a mixed linear model (MLM) and incorporated principal component analysis (PCA) with three components to account for population structure. Single nucleotide polymorphism (SNP) data from the 3k-chickpea genome sequencing project, along with phenotypic data, were retrieved from Varshney et al. (2021). Five major traits—days to flowering (DTF), days to maturity (DTM), hundred seed weight (HSW), plant height (PH), and pods per plant (PPP)—were selected to test the hypothesis. For each trait, the ‘GenoPlast score and mean’ were calculated, resulting in a total of 10 traits analyzed in the GWAS. We analyzed a dataset of 3,171 individuals using several stringent filters to ensure data quality and focus on common variants. We included only those SNPs present in at least 2,854 individuals, with minor allele frequencies ranging from 0.05 to 1.0, and limited the heterozygosity proportion to between 0.0 and 0.1. Additionally, we removed minor SNP states and restricted our analysis to biallelic SNPs to reduce complexity and potential sources of error. These criteria were applied to enhance the robustness of our findings by concentrating on well-genotyped, common genetic variants.

In the GWAS of 10 traits, significant marker-trait associations were detected for 5 traits after correcting for Bonferroni multiple correction. These traits are DTFm, DTFp, DTMm, HSWp, and HSWm, where ‘p’ stands for plasticity and ‘m’ stands for mean. The number of associated SNPs for each trait is as follows: DTFm with 24 SNPs, DTFp with 11 SNPs, DTMm with 14 SNPs, HSWp with 9 SNPs, and HSWm with 24 SNPs (Fig. 1A, Fig. S1). Overall, 60 SNPs were analyzed in total. The analysis revealed that the majority of SNPs (45 out of 60) were unique to individual traits, indicating distinct genetic associations for each trait. Additionally, 8 SNPs were shared between two traits, and 7 SNPs were shared among three traits, suggesting some level of pleiotropy (Fig. 1B). Notably, the traits related to plasticity (DTFp and HSWp) and mean (DTFm, DTMm, and HSWm) showed distinct SNP associations, highlighting the genetic differences between trait means and their plasticity. A chi-square test for independence between SNPs and traits yielded a chi-square statistic of 288.25 with a highly significant p-value of 6.89 ×10^−52^, indicating a strong association between SNPs and specific traits. This significant association reinforces the distinct genetic architecture underlying trait means and plasticity, suggesting that different sets of genetic factors contribute to the expression of these traits.

**Figure 1:**
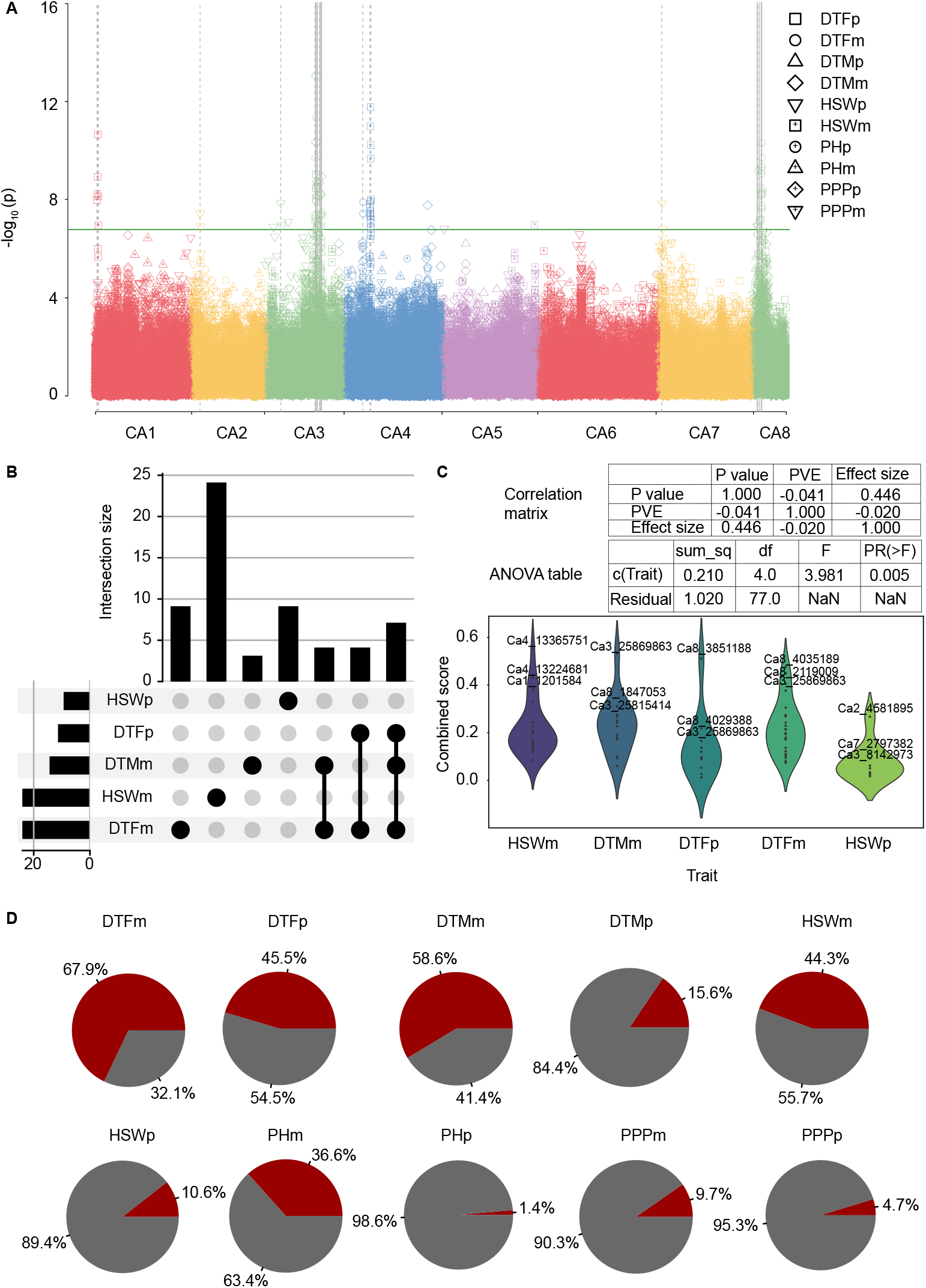
GWAS for 10 different traits in a 3k-panel of chickpea. A: The Manhattan plot displays the SNPs used for the GWAS analysis. The solid horizontal line represents the significance threshold after correction for the Bonferroni multiple testing. Solid vertical lines indicate instances where the same SNP is associated with the more than two traits, while dashed vertical lines indicate cases where the same SNP is associated with two different traits. B: An UpSet plot showing the significant SNPs that are unique to or shared among different traits, along with the number of SNPs in each category. C: A combined score calculated using p-values, phenotypic variance explained by the SNP, and effect size of the SNP. The violin plot illustrates the distribution of SNPs based on this combined score for the significant SNPs. D: The proportion of genetic versus residual contribution to traits identified in the GWAS analysis. Red indicates genetic contribution, while grey represents residual contribution.

To prioritize SNPs that are not only statistically significant but also biologically relevant, we used a combined score that integrates multiple metrics: p-value, effect size, and the proportion of phenotypic variance explained (PVE). By normalizing these metrics to a 0-1 scale, we ensured a standardized comparison and assigned weights of 0.4 to both p-value and PVE, and 0.2 to effect size, reflecting their relative importance. The correlation analysis revealed a moderate positive correlation between normalized p-values and effect sizes (0.446), but very low correlations between PVE and the other metrics, indicating their independence. ANOVA results showed a significant effect of traits on the combined score (*F* = 3.981, *p* = 0.005), and the Kruskal-Wallis test confirmed significant differences in combined scores across traits (*p* = 0.000441), underscoring the distinct genetic architectures for different traits (Fig. 1C).

Our analysis of the genetic and residual variances for the trait mean and plasticity across five traits (DTF, DTM, HSW, PH, and PPP) reveals distinct patterns. The mean genetic variance is consistently higher than the plasticity genetic variance for all traits, with DTF showing the highest mean genetic variance at 67.9% and PPP the lowest at 9.7%. Similarly, DTF has the highest plasticity genetic variance at 45.5%, while PH has the lowest at 1.4%. Conversely, the residual variance shows an inverse pattern, where the plasticity residual variance is generally higher than the mean residual variance. PPP exhibits the highest mean residual variance at 90.3%, and DTF the lowest at 32.1%. For plasticity, PH has the highest residual variance at 98.6%, while DTF has the lowest at 54.5% (Fig. 1D). The paired t-tests show significant differences in genetic (*p* = 0.0134) and residual variances (*p* = 0.0134) between the trait mean and plasticity. Overall, these patterns confirm that plasticity has a distinct genetic basis and can be considered a separate trait influenced by both genetic and environmental factors.

Our GWAS results provide evidence for the notion that the capacity for plastic response and the average expression of traits can evolve independently due to separate genetic controls. For example, the distinct SNP associations for DTFp and HSWp compared to DTFm and HSWm illustrate that plasticity in flowering time and seed weight is controlled by different genetic mechanisms than their respective mean traits (Fig. 1). These results help dispel the common misconception that plasticity, understood as a non-genetic mechanism, serves as a means of coping with environmental changes. We recognize plasticity itself as a distinct trait, analogous to other traits characterized by genetic polymorphisms. Plasticity emerges not solely from gene-environment interactions influencing the trait mean but also because the genes regulating plasticity are influenced by the same intricate interactions as other trait-related genes. These interactions encompass gene-environment interactions, epistasis, and pleiotropy. As a result, genetically diverse individuals may display varied phenotypic responses to identical environmental conditions due to differences in these plasticity genes.

Furthermore, GWAS analysis demonstrated that plasticity traits (DTFp and HSWp) had lower genetic variance compared to trait means (DTFm, DTMm, and HSWm), indicating lower heritability for plasticity. This suggests that environmental factors play a more significant role in shaping plasticity, leading to specialized individuals evolving under specific conditions. The higher genetic variance in trait means implies that these traits are under stronger genetic control, favoring stable trait expression in consistent environments. These findings align with the expectation that low heritability of plasticity leads to specialization, while higher heritability would favor increased plasticity and adaptability in individuals (Scheiner and Lyman, 1989).

In conclusion, our study underscores the importance of recognizing phenotypic plasticity as a distinct trait with its own genetic architecture. Understanding the genetic basis of plasticity can significantly enhance crop adaptation strategies in the face of rapid climate change, enabling the development of varieties that are both high-yielding and resilient across diverse environmental conditions.

## Acknowledgements

This work received support from the Department of Biotechnology, India, under Grant No. BT/PR23613/BAP /118/354/2017, awarded to E.R. A.G. is the recipient of an IISER Tirupati Institutional PhD Fellowship.

## Author contributions

Ganesh Alagarasan: Conceptualization; data curation; formal analysis; visualization; writing – original draft; writing – review and editing. Rajeev K. Varshney: chickpea GWAS project; writing – review and editing. Eswarayya Ramireddy: Conceptualization; methodology; supervision; writing – review and editing.

---

**Figure S1:**
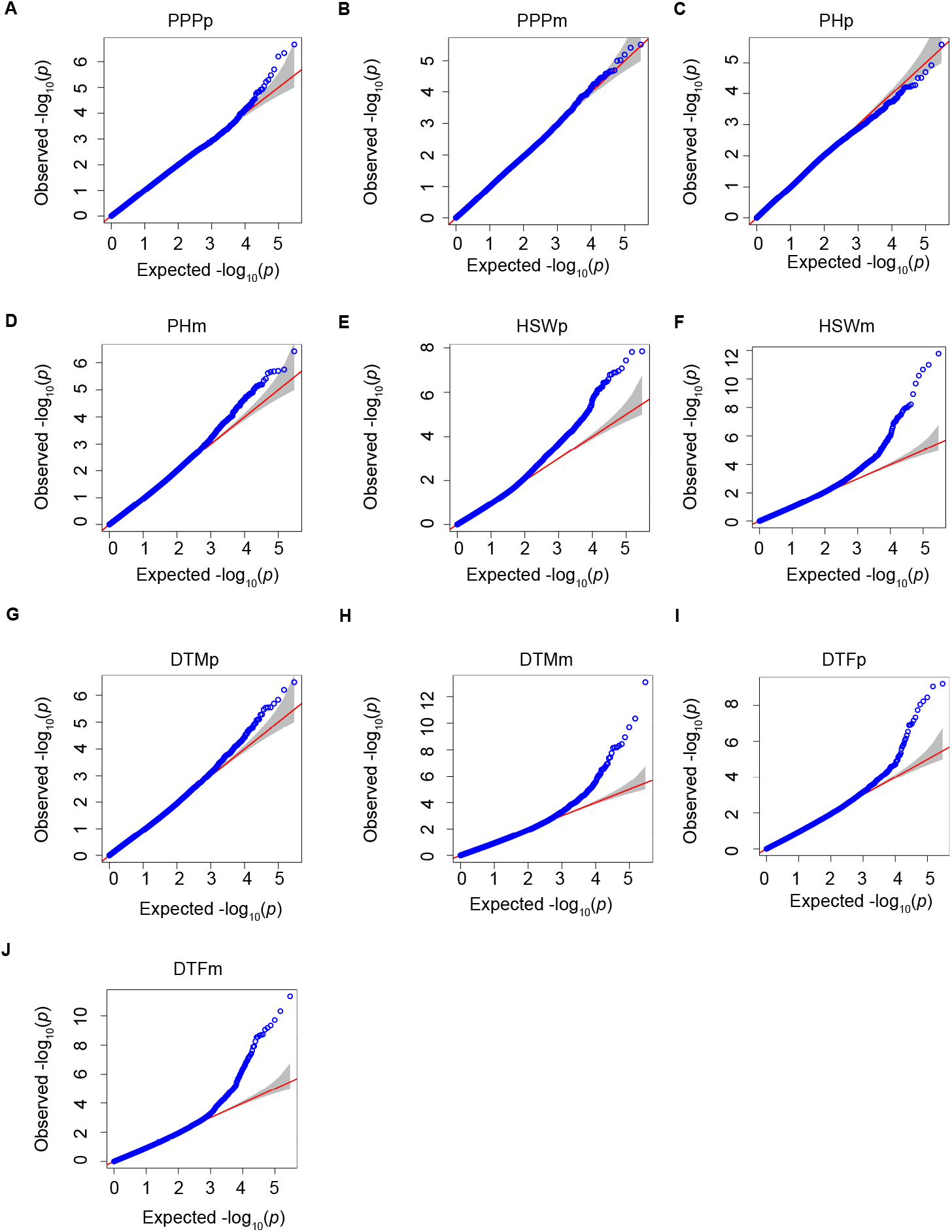
Quantile-Quantile (Q-Q) Plot of GWAS P-values. The Q-Q plot compares the observed p-values from the GWAS to the expected p-values under the null hypothesis of no association. The diagonal line represents the expected distribution under the null hypothesis. Points above the line indicate observed p-values lower than expected, suggesting potential genetic associations.

